# Denitrification of water in a microbial fuel cell (MFC) using seawater bacteria

**DOI:** 10.1101/107904

**Authors:** Samrat MVV Naga, Rao K Kesava, Bernardo Ruggeri, Tonia Tommasi

## Abstract

The sea contains various microbes which have an ability to reduce and oxidize substances like iron, sulphur, and nitrate. Most of these processes happen in the seawater, but can also be applied for purification of wastewater. In the present work, a consortium of seawater bacteria has been used for the first time in a microbial fuel cell to reduce nitrate in synthetic water samples and produce electricity by oxidizing organic matter. The concentrations of 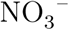 and 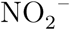 were reduced to well below their permissible limits. Moreover, the growth of the bacterial consortium at cathode causes an increased electricity production in the cell because of the increased bacterial activity. The performance of the cell with a bicarbonate buffered solution (BBS) at the cathode was superior to that obtained with the commonly used phosphate buffered solution (PBS). As BBS is the natural buffering agent found in the sea, the use of BBS is eco-friendly. The same seawater bacterial consortium could be used at both the anode and the cathode, confirming their adaptability to different environments. Unfortunately, denitrification was accompanied by the generation of high concentrations of 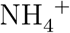 at the anode and the cathode, probably because of the use of N_2_ gas for sparging the anolyte. This aspect merits further investigation.

## 1. Introduction

Compounds containing nitrogen are abundant, with an annual production of about 4.13 x 10^14^ g as N by fixation processes (Zhang and Zindler, 1993). Among them, nitrate 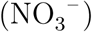 is one of the predominant compounds produced which is a highly mobile and stable species. Also, most of the compounds are converted to 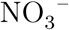 after their use in biological systems (Galloway, 1998). The released 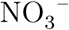 enters the groundwater sources in the form of domestic and industrial wastewater, and fertilizers that are added to the soil for higher crop yields. This leads to widespread environmental contamination (Kundu and Mandal, 2009; Zhang et al., 2015; Lasagna et al., 2016; Nakagawa et al., 2016).

When nitrate is ingested by people, it is converted to nitrite and then to nitrosamines, which may cause gastric cancer (Bogárdi et al., 2013). Nitrate intake causes methaemoglobinaemia in infants, even when exposed for a very short duration (Fewtrell, 2004). The World Health Organization (WHO) has prescribed guideline values (GV) of 50 mg 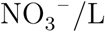 and 10 mg 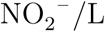 (World Health Organization, 2011). To obtain concentrations less than the GV, methods such as adsorption, reverse osmosis, ion exchange, electrodialysis, catalytic denitrification, and biological denitrification have been used (Hell et al., 1998; Schoeman and Steyn, 2003; Samatya et al., 2006; Ghafari et al., 2008; Bhatnagar and Sillanpää, 2011).

Because of the disadvantages of many of the processes, it is preferable to use the biological route, wherein bacteria can reduce 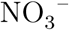 to N_2_ gas by donating the electrons which are released in their metabolic pathways (Munn, 2011). An additional disinfection step such as UV treatment or passing the water through a bed of silver nanoparticles (Hijnen et al., 2006; Swathy et al., 2014) is needed before the water can be used for drinking.

The reduction and oxidation steps can be made to occur in the anodic and cathodic compartments of a microbial fuel cell. This is similar to a conventional fuel cell, where the oxidation of substrates occurs at the anode, and reduction occurs at the cathode. When oxidation-reduction and transfer of electrons occur in the presence of, or are mediated by microorganisms, then it is called a microbial fuel cell (MFC). The current obtained is produced by the oxidation of organic substrates and the produced electrons (e^−^) are used for the reduction of the nitrate at the cathode (Fig. 1).

**Figure 1:**
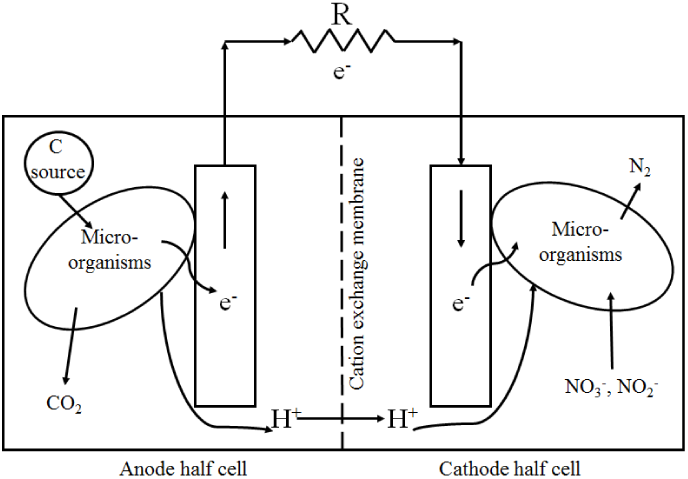
Schematic of a microbial fuel cell. Here R represents a resistor.

The overall reaction is (Zumft, 1997; Clauwaert et al., 2007)

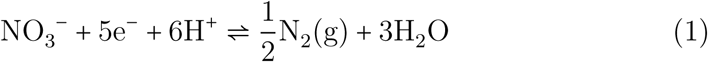

The reduction of 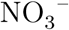 mostly occurs in the absence of O_2_, as the enzymes involed in the denitrification process are repressed by O_2_. Therefore an anaerobic condition is maintained in the cell (Munn, 2011). The MFC produces nitrate-free water and also generates electricity. The first application of this method for nitrate removal was by Clauwaert et al. (2007), who observed complete denitrification with power and current densities of about 4 W/m^3^ TCC and 19 A/m^3^ TCC, respectively, and a cell voltage of 0.214 V. Here TCC denotes the total cathodic compartment volume. Similar experiments, but with simultaneous carbon, nitrate, or phenol removal were performed with different COD/N (chemical oxygen demand/nitrogen) ratios (Virdis et al., 2008, 2010; Zhang and He, 2012; Feng et al., 2015).

Attempts were made to reduce the resistances at the cathode side by adjusting the pH, using transient operation, and by design modifications (Clauwaert et al., 2009; Behera et al., 2010; Liang et al., 2013; Zhu et al., 2013; Yang et al., 2015). In most of these systems, phosphate buffered solutions (PBS) were used as the media to control the pH and also to provide sufficient conductivity for the flow of ions. The increased use of phosphate and its subsequent discharge into water sources causes eutrophication, which is a serious environmental problem (Morse et al., 1998; Loganathan et al., 2014). Therefore to limit the use of PBS, buffers such as bicarbonate, and boric acid-borate were used (Fan et al., 2007; Chen et al., 2015).The power density at the anode with a bicarbonate buffered solution (BBS) was about 39% higher compared to PBS (Fan et al., 2007).

There are also reports on the direct use of low-conductivity waters like groundwater sources without buffer addition, and with a high nitrate content, and their subsequent denitrification. Pous et al. (2013) observed a 64% removal efficiency of nitrate for a 15-day period of operation using water with a conductivity less than 1000 *µ*S/cm. For water with higher conductivities in the range 1000 - 4000 *µ*S/cm, Puig et al. (2012) observed about 45 - 90% removal when the system was operated for 3 days. Groundwater sources were also treated using PBS, but with a modified design of the fuel cell. Anion exchange membranes were used to permit the movement of anions from the surrounding groundwater to the buffered anolyte and catholyte solutions, which results in desalination and denitrification (Tong and He, 2013; Zhang and Angelidaki, 2013). As bicarbonate is a naturally occurring buffering compound (Weber and Stumm, 1963), its use instead of phosphate can help in reducing the load of phosphate contamination on the environment.

The methods discussed above for denitrification either used a single strain of bacteria, or a consortium taken from a waste sludge or a bio reactor used in wastewater treatment plants. Denitrifying and nitrifying bacteria are also present to a greater extent in seawater. It is estimated that annually about 1.4 x 10^14^ g N is fixed by marine ecosystems and about 1.0 x 10^14^ − 2.8 x 10^14^ g N is denitrified to N_2_ gas in the oceans (Fowler et al., 2013). Therefore, the use of seawater bacteria as an inoculum may increase the rates of denitrification. In the present work, seawater bacteria taken from Adriatic sea at the shores of Italy have been used. There are reports on the use of seawater bacteria for oxidizing substrates at the anode of a MFC (Tommasi et al., 2016), but there are no reports on their use for denitrification.

The nitrate removal capability of these bacteria was examined using PBS and BBS at the cathode. Linear sweep voltammetry measurements were made to confirm the superiority of BBS compared to PBS. The present work is mainly confined to the results obtained with BBS.

## 2. Materials and methods

### 2.1. Experimental setup and materials

Two different microbial fuel cell (MFC) designs were used in the experiments. Both the designs consist of two chambers, an anode and a cathode, and were made of poly(methyl methacrylate). The dimensions of a single rectangular and circular chamber were 8 x 8 x 2 cm^3^ (volume ≈ 128 mL), and 7 cm (diameter) and 15 mm (width) (volume ≈ 57.7 mL), respectively. All the experiments were conducted in duplicate under the same operating conditions. The anode and cathode chambers were separated by a cation exchange membrane (CEM) to facilitate the movement of cations from the cathode to the anode and vice-versa. For the movement of electrons from the anode to cathode through the external circuit, graphite rods were stitched to a commercial carbon felt (C-FELT) (soft felt SIGRATHERM GFA 5, SGL Carbon, Germany) using a conductive carbon thread and the rods were connected to a potentiostat using copper wires with crocodile clips. The carbon felt was used to increase the effective area exposed to increase the electron transfer.

The rectangular cell was operated in a semi-batch mode, wherein an initial amount of substrate was added to the chamber. Sampling and addition of new substrate was done periodically. The circular shape MFC was operated in a semi-continuous mode, wherein the solution was re-circulated from the chamber to reservoir using a peristaltic pump (ISMATEC-ISM404B, Germany) at a flow rate of 36 mL/min (Fig. 2). During oxidation and reduction, gases are released at the electrodes. The gas released in the chamber, and the liquid enter the bottle A, where the gas is separated and enters the bottle B. The water vapour associated with the humid gas is condensed in B, and the relatively dried gas enters the bottle C, which contains water at a pH = 2.0. The low pH prevents the dissolution of CO_2_. Hence the gas displaces the water into the bottle D (Fig. 2). The volume of water collected in D is equal to the volume of gas released from the chamber. The total volume of the solution taken was about 110 mL in rectangular chamber and about 200 mL in the circular chamber and the bottle A.

**Figure 2:**
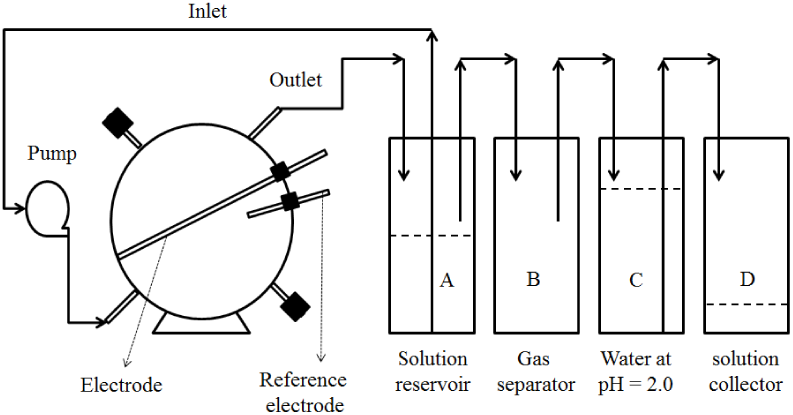
Schematic of the circular MFC setup used in the experiments.

The chemicals sodium acetate (AR), peptone (bacteriological for microbiology), sodium phosphate dibasic dihydrate (AR), sodium phosphate monobasic monohydrate (AR), potassium nitrate (AR), sodium bicarbonate (AR) were purchased from Sigma Aldrich and used to prepare the anolyte and catholyte without further purification. Ultrapure Millipore water was used to prepare anolyte solution. For the catholyte, commercially and locally available mineral bottled water was used.

### 2.2. MFC startup and operation

The MFCs used in the experiments were inoculated with the effluent from an active MFC that was previously enriched. The enrichment procedure is given in Tommasi et al. (2016), who also present a detailed discussion of the different types of bacteria in the seawater consortia. The consortium consisted of species such as Proteobacteria phylum (*Shewanella* and *Geobacter*), Firmicutes (*Clostridium*) and Ascomycota (*Saccharomyces*). Phosphate and bicarbonate buffers of approximately 80 mM concentration were prepared. Phosphate buffer was prepared by the addition of 8.2 g/L Na_2_HPO_4_ ⋅ 2 H_2_O and 5.9 g/L NaH_2_PO_4_ ⋅ 2 H_2_O to distilled water. Bicarbonate buffer was prepared by the addition of 5.46 g/L NaHCO_3_, 48.1 mg/L Na_2_HPO_4_ ⋅ 2 H_2_O and 35.9 mg/L NaH_2_PO_4_ ⋅ 2 H_2_O to mineral water. The anolyte used consisted of PBS with 10 mL inoculum, 8 g/L CH_3_COONa, 10 g/L peptone, 12 g/L C_6_H_12_O_6_. As we are interested in denitrification of water, a catholyte containing 0.16 g/L KNO_3_ (100 mg NO_3_/L) in BBS was used. To avoid substrate limitations at the anode for the production of e^−^, the anolyte had a higher concentration of the carbon source than that required by stoichiometry. All the solutions were adjusted to a pH between 7 - 7.5 before their injection into the MFC using 2 N NaOH or HCl solution. Phosphate buffered solution and BBS were purged with nitrogen and argon gas, respectively for about 5 minutes, to maintain the cells in an anaerobic condition. The solution volumes in the anode and the cathode chambers of the cells were kept constant. The samples from the chambers were collected periodically and an equal amount of fresh solution was added.

A chamber containing a cathode, an electrolyte and bacteria will be referred to as a biotic cathode, and one without bacteria will be referred to as an abiotic cathode. Similar remarks apply to the anode. In the present work, a biotic anode was used along with abiotic and biotic cathodes.

The solution containing bacteria was prepared by adding 1 mL of the enriched inoculum to 10 mL of the solution. Initial studies were done using rectangular cell with an abiotic cathode but with biotic mode in anode. To confirm the dual nature of seawater bacteria, the bacteria enriched in the anodic compartment during the experiments with an abiotic cathode were used in the cathodic compartment for denitrification. The biotic experiments at cathode were performed using the circular setup for BBS to decrease the limitations that can arise because of slow diffusion of species compared to rectangular setup with PBS.

All the cells used were initially connected under open circuit voltage to attain a steady voltage. Then the cells were connected to a resistance of 1.5 kΩ. Periodic potentiostat measurements was conducted using linear sweep voltammetry (LSV).

### 2.3. Electrochemical measurements on MFCs

The electrochemical experiments were conducted using a multi-channel VSP potentiostat/galvanostat (BioLogic, France). The data from these experiments were acquired using a EC-Lab software version 10.1x. Polarization curves i.e. the variation of the cell potential with current density, were obtained using LSV by imposing a linear decrease of the electric potential from the open circuit voltage (OCV) (current *I* = 0) to the short circuit voltage (SCV) (*I* = *I*_*max*_) of the cell, at a scan rate of 1 mV/s. From these *I* − *V* curves, the power density and the current density were calculated using the equations *I*_*d*_ = *I/V*_*c*_ and *P*_*d*_ = (*IV*_*o*_)/*V*_*c*_, where *I*_*d*_ and *P*_*d*_ are current and power density, respectively, and *V*_*o*_ and *V*_*c*_ represent the voltage output and the volume of the cathodic chamber, respectively. These measurements were done for a full cell, and also for the individual electrodes. For a full cell, a two-electrode configuration was used, where the working electrode (WE) was connected to the anode and both the counter (CE) and the reference (RE) electrodes were connected to the cathode.

### 2.4. Chemical analysis of samples

Samples collected at certain time intervals were analysed using a Metrohmion chromatograph for anions 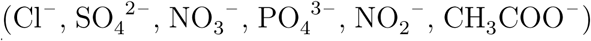 and cations 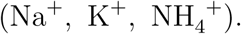 When PBS was used, the eluent for the anions contained 2.5 mM Na_2_CO_3_, 1.0 mM NaHCO_3_, and 2.0 mM NaOH. The samples were diluted about 200 times before injection into the column. When BBS was used, the eluent for the anions contained 1.0 mM NaHCO_3_ and 3.2 mM Na_2_CO_3_. The samples were diluted 25 times. The separation of ions was achieved using a Metrosep A Supp 5 - 250/4.0 anion column with a Metrosep Guard column. The cation analysis was done using a Metrosep C4 - 150/4.0 column. The eluent used for the cations was a mixture of 1.7 mM HNO_3_ and 0.7 mM dipicolinic acid. The samples were diluted 200 times before injection into column.

The composition of the gaseous samples was determined using an offline gas chromatograph (VARIAN CP4900), where H_2_, N_2_, CO_2_, and O_2_ were detected.

## 3. Results and discussion

### 3.1. Ability of seawater bacteria to denitrify water

Denitrification occurred at a faster rate with a biotic cathode (i.e. one having seawater bacteria) than an abiotic cathode (Fig. 3). In Figs. 3 - 5, 8, and S2, the error bars represent the 95% confidence intervals. At the end of two days, the removal of nitrate was about 100% and 6% with the biotic and abiotic cathodes, respectively. Also, nitrite was not detected in the system. Thus there was a total conversion of nitrate to N_2_. This was also confirmed by the gas analysis of the samples collected, which contained about 99.8% N_2_ gas. Thus, a consortium of bacteria taken from the interface between the sea and the air, where most of the nitrogen-fixing bacteria are present (Zhang and Zindler, 1993), significantly enhanced the rate of denitrification.

**Figure 3:**
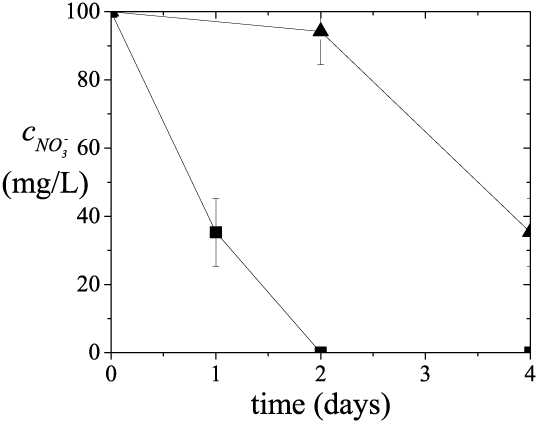
Variation of the concentration of nitrate in the catholyte with time: ▲, abiotic cathode or one without bacteria; ∎, biotic cathode. A PBS was used at the cathode of the rectangular cell.

The use of PBS to examine the denitrification capacity of bacteria is good on a lab scale. However, it is not feasible at the field level, as the presence of high concentrations of phosphates in water causes algal growth, and the use of such water for drinking is not advisable. In the case of seawater bacteria, the use of bicarbonate is a sustainable solution for denitrification, as it is a natural buffering agent present in the sea. Also, it mimics the environmental conditions to which the seawater bacteria are acclimatized. Nitrate and nitrite levels very much lower than GV values prescribed by WHO (50 mg/L for 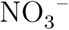 and 10 mg/L for 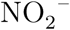 were obtained when BBS was used at the cathode (Fig. 4a).

**Figure 4:**
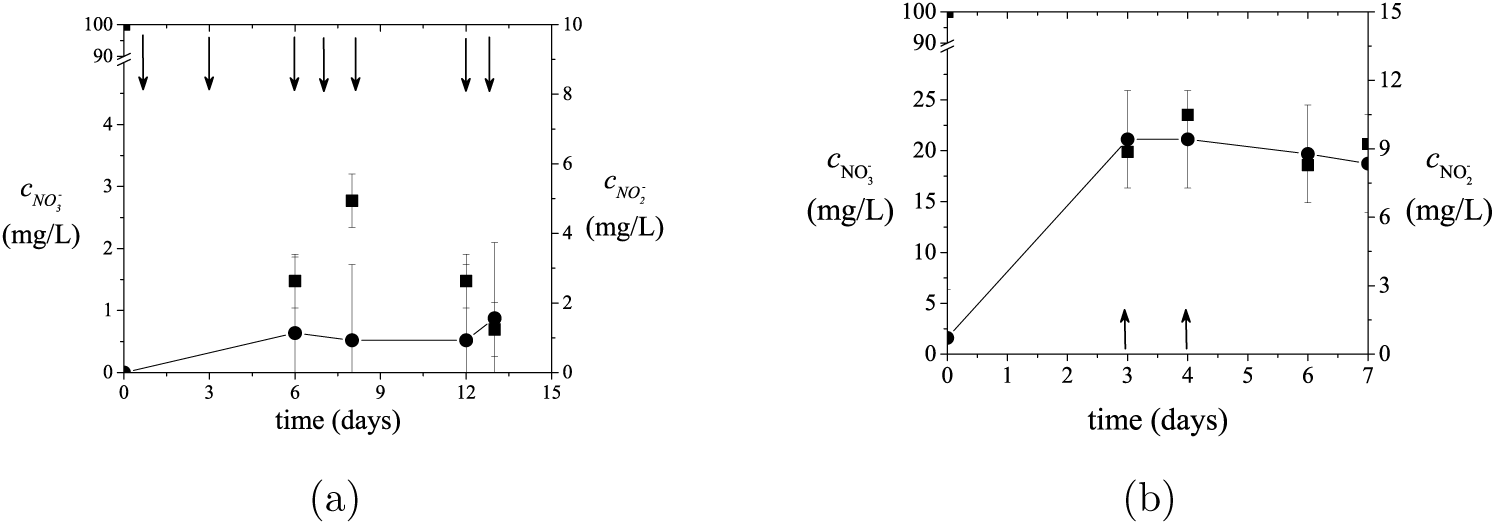
Variation of the concentration of nitrate, ∎ and nitrite, ● in the biotic cathode with time: concentration of BBS at the cathode = 80 mM (a), 160 mM (b). The arrows represent an injection of 10 mL of fresh catholyte. The circular cell was used at the cathode.

The nitrate reduction at the biotic cathode is mostly dependent on the availability of the electrons at the cathode and also on the mass transfer of the nitrate from the bulk solution to the cathode surface. In order to reduce this resistance, the catholyte was recirculated. Also, to permit free movement of the ions, the conductivity of the catholyte was increased by doubling its concentration to 160 mM (conductivity ≈ 9.8 mS/cm). When the concentration of the bicarbonate was increased, there was a 9-fold increase in the concentration of 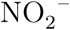, and a decrease in the amount of 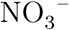 reduced (Fig. 4b). Such a decrease of the denitrification capability of bacteria with an increase of conductivity has not been reported earlier. However, there are reports of the decrease in performance as the conductivity was increased for experiments with a bio-anode and a bio-cathode using an aerobic process (De Schamphelaire et al., 2010; Lefebvre et al., 2012; Karthikeyan et al., 2016). It has been reported that the presence of higher salt concentrations can adversely affect the physiology of bacteria, as the salt tolerance for different organisms is different. This may explain the ability of seawater bacteria to denitrify solutions of lower conductivity compared to those of higher conductivity. The electrons needed for the reduction of nitrate are liberated by the oxidation of glucose and acetate at the anode. The concentration of acetate in the anode compartment decreases with time, except for a sudden increase when fresh anolyte is added (Fig. 5).

**Figure 5:**
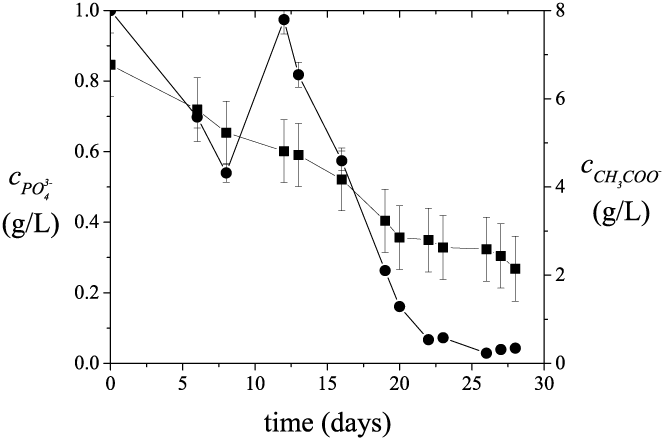
Variation of the concentration of phosphate in the biotic catholyte, ∎ and acetate at the biotic anolyte, ● with time for the circular cell. Here the cathode and anode contain BBS and PBS, respectively. For reasons discussed in the text, the BBS contains some phosphate.

There was a steady decrease in the concentration of phosphate in the cathode compartment (Fig. 5), probably because of the use of phosphate as a nutrient source by the bacteria. Here the presence of phosphate in BBS is mainly because of the addition of inoculum from a previous MFC which was operated with PBS. An increase of the optical density of the catholyte confirms the growth of bacteria (Supplementary Fig. S1). This indicates that the bacteria used nitrate, phosphate, and the electrons from the anode for their growth. Hence, there was a stable removal of nitrate from the solution (Fig. 4).

### 3.2. Higher power generation with BBS at the cathode

The reduction of the nitrate at the cathode is possible because of the availability of the electrons produced at the anode. Open circuit voltage (OCV) condition were initially used for 1 day and 5 days for the rectangular and the circular cells, respectively, to get the bacterial population acclimatized to the conditions in the cell. When PBS was used at the biotic cathode, the open circuit voltage (*V*_*OCV*_) was about 0.37 V. When the BBS was used as the catholyte in the circular cell, there was an increase in *V*_*OCV*_ to 0.45 V (Fig. 6). There was also significant increase in the maximum power and current densities from 0.6 W/m^3^ and 7.5 A/m^3^ to 2.1 W/m^3^ and 26.6 A/m^3^, respectively, as the buffer solution was changed (Fig. 6).

**Figure 6:**
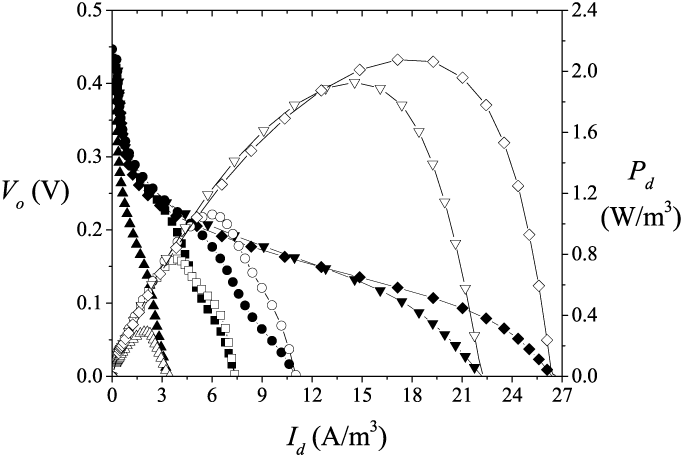
Polarization (filled symbols) and power density (open symbols) curves for the circular cell with BBS at the cathode and measured at different times during the course of the test: ∎, ◻, 6 days; ●,○, 6 days, after replenishment; ▲, △, 8 days; ▼, ▽, 12 days, after replenishment; ◆,⋄, 14 days.

There was a sudden decrease in the voltage as the current density was increased. This is caused by charge transfer mechanisms (Zhao et al., 2009). The OCV was almost constant with respect to time, but there was a considerable increase in the peak power and current densities after 8 days (Fig. 6). This is mainly because of the availability of sufficient substrate for the oxidation and reduction reactions to occur and generate electrons.

The anode was buffered using PBS having a conductivity of ≈ 10 mS/cm, which is 40% higher than that of BBS. Therefore, to check the performance of the cell with a higher BBS conductivity at the cathode, the concentration of bicarbonate was doubled. There was an almost two-fold increase in the current and the power density, even though there was very less change in the OCV (Fig. 7). As mentioned earlier, with an increase in the bicarbonate concentration, there was a decrease in the reduction of 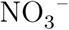 and 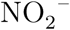 to N_2_ (Fig. 4). However, based on the measurement of the current density, there is an extra flow of electrons from the anode to the cathode because of the increased conductivity. This should cause an increased reduction of 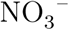 contrary to observation. It is possible that the N_2_ gas produced may further reduced to 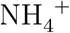 by the reaction (Kim et al., 2005).

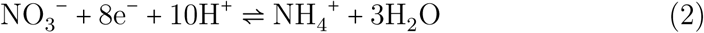

**Figure 7:**
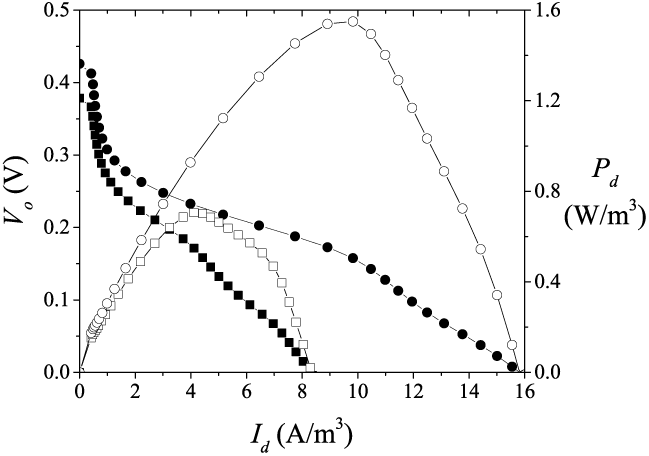
Polarization (filled symbols) and power density (open symbols) curves for the full cell with BBS at the cathode, measured before (∎, ◻) and after (●,○) addition of the concentrated bicarbonate solution.

The competition for e^−^ by the reactions (1) and (2) may cause a decrease in the reduction of 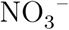.

### 3.3. Biological denitrification and a possible chemical nitrification in the cell

The reduction of nitrate with the use of electrons and protons occurs as shown by the reaction (1). With an excess availability of electrons it is possible to reduce the nitrate and nitrite to 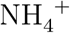 by the reactions (2) and (3) (Kim et al., 2005)

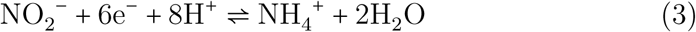

In our experiments, 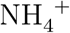 ions were produced at the cathode and the anode (Fig. 8). There was a continuous movement of H^+^ from the anode to the cathode to compensate for the e^−^ that passed to the cathode through the external circuit. The 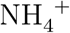 ion may move to the anode to account for the loss of H^+^. Perhaps 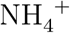 is the only compensating ion as there was a negligible change in the concentration of Na^+^, which is the major cation in the system (Supplementary Fig. S2). This increased the concentration of 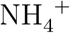 at the anode (Fig. 8b). When the concentration of bicarbonate was doubled, there was an adjustment in the concentrations of Na^+^ to attain a constant value. The concentration of 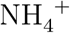 in the anode was very high compared to its value at the cathode. Thus 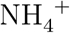 appears to diffuse against its concentration gradient, a phenomenon that has been observed in multicomponent systems (Krishna, 2016). However, in view of the large difference in concentrations, further investigation is needed. As the solution at the anode was initially sparged with N_2_ gas, dissolved N_2_ can diffuse from the anode to the cathode. With the availability of excess e^−^ and H^+^, N_2_ may be reduced to 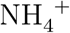 at the cathode, which may diffuse to the anode.

**Figure 8:**
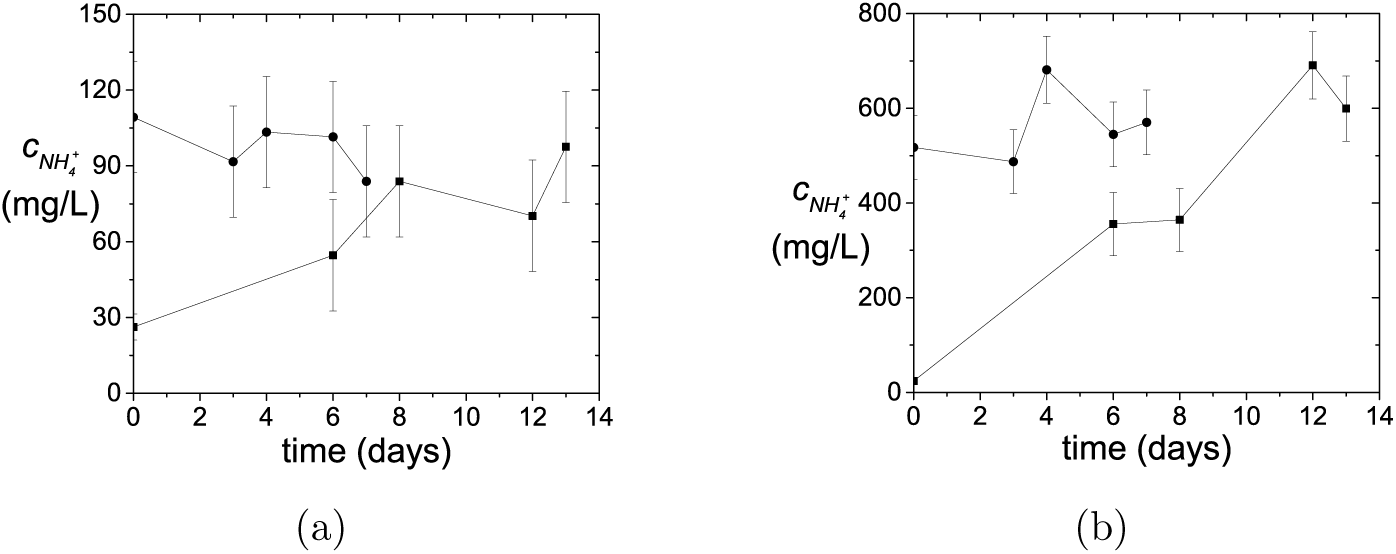
Variation of the concentration of 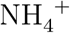 with time in the (a) catholyte and (b) anolyte: ∎, 80 mM 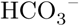 ●, 160 mM 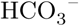 at the cathode.

If a bacterial population has to achieve nitrification at the anode using a species such as *Planctomycetes* (anammox bacteria), it requires 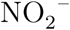 with 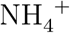 for the oxidation reaction (Kuenen, 2008)

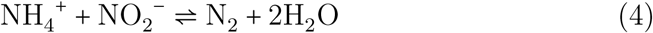

As there is no nitrite present at the anode, the only reaction that can occur is (Kim et al., 2005, 2006)

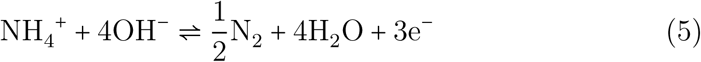

. Thus chemical nitrification occurs at the anode and biological denitrification occurs at the cathode.

## 4. Conclusions

Seawater bacteria could reduce 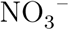 to N_2_, and the levels of 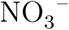 and 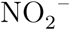 were well below the guideline values prescribed by the World Health Organization. The bacteria were able to function in a phosphate buffered solution (PBS) at the anode and bicarbonate buffered solution (BBS) at the cathode. The BBS is environmentally friendly and can denitrify water with higher power and current densities in the presence of seawater bacteria. However, a high concentration of 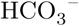 (160 mM) decreased the denitrification capacity of the cell. Hence, seawater bacteria can be used for the denitrification of water with low to medium concentrations of bicarbonate. The high concentrations of 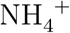 at the cathode and the anode may result from sparging the anolyte with N_2_ or because of the presence of higher amounts of substrate at anode which results in the production of higher number of e^−^. It appears that chemical nitrification occurs at the anode. Therefore, a proper selection of the sparging gas and the use of a stoichiometric amount of substrate is necessary.

## Acknowledgements

NS would like to thank the European Commission for providing the Erasmus Mundus fellowship to conduct the research at the POLITO.

## References

Behera, M., Jana, P. S., Ghangrekar, M., 2010. Performance evaluation of low cost microbial fuel cell fabricated using earthen pot with biotic and abiotic cathode. Bioresource Technol. 101, 1183–1189.

Bhatnagar, A., Sillanpää, M., 2011. A review of emerging adsorbents for nitrate removal from water. Chem. Eng. J. 168, 493–504.

Bogárdi, I., Kuzelka, R. D., Ennenga, W., 2013. Nitrate Contamination: Exposure, Consequence, and Control. Vol. 30. Springer Science & Business Media.

Chen, G., Zhang, S., Li, M., Wei, Y., 2015. Simultaneous pollutant removal and electricity generation in denitrifying microbial fuel cell with boric acid-borate buffer solution. Water Sci. Technol. 71, 783–788.

Clauwaert, P., Desloover, J., Shea, C., Nerenberg, R., Boon, N., Verstraete, W., 2009. Enhanced nitrogen removal in bio-electrochemical systems by pH control. Biotechnol. Lett. 31, 1537–1543.

Clauwaert, P., Rabaey, K., Aelterman, P., De Schamphelaire, L., Pham, T. H., Boeckx, P., Boon, N., Verstraete, W., 2007. Biological denitrification in microbial fuel cells. Environ. Sci. Technol. 41, 3354–3360.

De Schamphelaire, L., Boeckx, P., Verstraete, W., 2010. Evaluation of biocathodes in freshwater and brackish sediment microbial fuel cells. Appl. Microbiol. Biot. 87, 1675–1687.

Fan, Y., Hu, H., Liu, H., 2007. Sustainable power generation in microbial fuel cells using bicarbonate buffer and proton transfer mechanisms. Environ. Sci. Technol. 41, 8154–8158.

Feng, C., Huang, L., Yu, H., Yi, X., Wei, C., 2015. Simultaneous phenol removal, nitrification and denitrification using microbial fuel cell technology. Water Res. 76, 160–170.

Fewtrell, L., 2004. Drinking-water nitrate, methemoglobinemia, and global burden of disease: A discussion. Environ. Health Persp. 112, 1371–1374.

Fowler, D., Coyle, M., Skiba, U., Sutton, M. A., Cape, J. N., Reis, S., Sheppard, L. J., Jenkins, A., Grizzetti, B., Galloway, J. N., Vitousek, P., Leach, A., Bouwman, A. F., Butterbach-Bahl, K., Dentener, F., Stevenson, D., Amann, M., Voss, M., 2013. The global nitrogen cycle in the twenty-first century. Phil. Trans. R. Soc. B 368, 1–14.

Galloway, J. N., 1998. The global nitrogen cycle: Changes and consequences. Environ. Pollut. 102, 15–24.

Ghafari, S., Hasan, M., Aroua, M. K., 2008. Bio-electrochemical removal of nitrate from water and wastewater: A review. Bioresource Technol. 99, 3965–3974.

Hell, F., Lahnsteiner, J., Frischherz, H., Baumgartner, G., 1998. Experience with full-scale electrodialysis for nitrate and hardness removal. Desalination 117, 173–180.

Hijnen, W., Beerendonk, E., Medema, G. J., 2006. Inactivation credit of UV radiation for viruses, bacteria and protozoan (oo) cysts in water: A review. Water Res. 40, 3–22.

Karthikeyan, R., Selvam, A., Cheng, K. Y., Wong, J. W.-C., 2016. Influence of ionic conductivity in bioelectricity production from saline domestic sewage sludge in microbial fuel cells. Bioresource Technol. 200, 845–852.

Kim, K.-W., Kim, Y.-J., Kim, I.-T., Park, G.-I., Lee, E.-H., 2005. The electrolytic decomposition mechanism of ammonia to nitrogen at an IrO_2_ anode. Electrochim. Acta 50, 4356–4364.

Kim, K.-W., Kim, Y.-J., Kim, I.-T., Park, G.-I., Lee, E.-H., 2006. Electrochemical conversion characteristics of ammonia to nitrogen. Water Res. 40, 1431–1441.

Krishna, R., 2016. Highlighting coupling effects in ionic diffusion. Chem. Eng. Res. Des. 114, 1–12.

Kuenen, J. G., 2008. Anammox bacteria: from discovery to application. Nat. Rev. Microbiol. 6, 320–326.

Kundu, M. C., Mandal, B., 2009. Nitrate enrichment in groundwater from long-term intensive agriculture: Its mechanistic pathways and prediction through modeling. Environ. Sci. Technol. 43, 5837–5843.

Lasagna, M., De Luca, D. A., Franchino, E., 2016. Nitrate contamination of groundwater in the western Po plain (Italy): The effects of groundwater and surface water interactions. Environ. Earth Sci. 75, 1–16.

Lefebvre, O., Tan, Z., Kharkwal, S., Ng, H. Y., 2012. Effect of increasing anodic NaCl concentration on microbial fuel cell performance. Bioresource Technol. 112, 336–340.

Liang, P., Yuan, L., Wu, W., Yang, X., Huang, X., 2013. Enhanced performance of bio-cathode microbial fuel cells with the applying of transient-state operation modes. Bioresource Technol. 147, 228–233.

Loganathan, P., Vigneswaran, S., Kandasamy, J., Bolan, N. S., 2014. Removal and recovery of phosphate from water using sorption. Crit. Rev. Env. Sci. Tec. 44, 847–907.

Morse, G., Brett, S., Guy, J., Lester, J., 1998. Review: Phosphorus removal and recovery technologies. Sci. Total Environ. 212, 69–81.

Munn, C., 2011. Marine Microbiology. Garland Science.

Nakagawa, K., Amano, H., Asakura, H., Berndtsson, R., 2016. Spatial trends of nitrate pollution and groundwater chemistry in Shimabara, Nagasaki, Japan. Environ. Earth Sci. 75, 1–17.

Pous, N., Puig, S., Coma, M., Balaguer, M. D., Colprim, J., 2013. Bioremediation of nitrate-polluted groundwater in a microbial fuel cell. J. Chem. Technol. Biot. 88, 1690–1696.

Puig, S., Coma, M., Desloover, J., Boon, N., Colprim, J., Balaguer, M. D., 2012. Autotrophic denitrification in microbial fuel cells treating low ionic strength waters. Environ. Sci. Technol. 46, 2309–2315.

Samatya, S., Kabay, N., Yüksel, Ü., Arda, M., Yüksel, M., 2006. Removal of nitrate from aqueous solution by nitrate selective ion exchange resins. React. Funct. Polym. 66, 1206–1214.

Schoeman, J., Steyn, A., 2003. Nitrate removal with reverse osmosis in a rural area in South Africa. Desalination 155, 15–26.

Swathy, J., Sankar, M. U., Chaudhary, A., Aigal, S., Pradeep, T., 2014. Antimicrobial silver: An unprecedented anion effect. Sci. Rep. 4.

Tommasi, T., Sacco, A., Armato, C., Hidalgo, D., Millone, L., Sanginario, A., Tresso, E., Schilirò, T., Pirri, C. F., 2016. Dynamical analysis of microbial fuel cells based on planar and 3D-packed anodes. Chem. Eng. J. 288, 38–49.

Tong, Y., He, Z., 2013. Nitrate removal from groundwater driven by electricity generation and heterotrophic denitrification in a bioelectrochemical system. J. Hazard. Mater. 262, 614–619.

Virdis, B., Rabaey, K., Rozendal, R. A., Yuan, Z., Keller, J., 2010. Simultaneous nitrification, denitrification and carbon removal in microbial fuel cells. Water Res. 44, 2970–2980.

Virdis, B., Rabaey, K., Yuan, Z., Keller, J., 2008. Microbial fuel cells for simultaneous carbon and nitrogen removal. Water Res. 42, 3013–3024.

Weber, W. J., Stumm, W., 1963. Buffer system of natural fresh water. J. Chem. Eng. Data 8, 464–468.

World Health Organization, 2011. Guidelines for Drinking-Water Quality, 4th Edition. World Health Organization (WHO).

Yang, Q., Zhao, H., Liang, H., 2015. Denitrification of overlying water by microbial electrochemical snorkel. Bioresource Technol. 197, 512–514.

Zhang, F., He, Z., 2012. Integrated organic and nitrogen removal with electricity generation in a tubular dual-cathode microbial fuel cell. Process Biochem. 47, 2146–2151.

Zhang, Q., Sun, J., Liu, J., Huang, G., Lu, C., Zhang, Y., 2015. Driving mechanism and sources of groundwater nitrate contamination in the rapidly urbanized region of south China. J. Contam. Hydrol. 182, 221–230.

Zhang, Y., Angelidaki, I., 2013. A new method for in situ nitrate removal from groundwater using submerged microbial desalination–denitrification cell (SMDDC). Water Res. 47, 1827–1836.

Zhang, Y., Zindler, A., 1993. Distribution and evolution of carbon and nitrogen in earth. Earth Planet. Sc. Lett. 117, 331–345.

Zhao, F., Slade, R. C., Varcoe, J. R., 2009. Techniques for the study and development of microbial fuel cells: An electrochemical perspective. Chem. Soc. Rev. 38, 1926–1939.

Zhu, G., Onodera, T., Tandukar, M., Pavlostathis, S. G., 2013. Simultaneous carbon removal, denitrification and power generation in a membrane-less microbial fuel cell. Bioresource Technol. 146, 1–6.

Zumft, W. G., 1997. Cell biology and molecular basis of denitrification. Microbiol. Mol. Biol. R. 61, 533–616.

